# Sequence determinants of protein phase behavior from a coarse-grained model

**DOI:** 10.1101/238170

**Authors:** Gregory L. Dignon, Wenwei Zheng, Young C. Kim, Robert B. Best, Jeetain Mittal

## Abstract

Membraneless organelles important to intracellular compartmentalization have recently been shown to comprise assemblies of proteins which undergo liquid-liquid phase separation (LLPS). However, many proteins involved in this phase separation are at least partially disordered. The molecular mechanism and the sequence determinants of this process are challenging to determine experimentally owing to the disordered nature of the assemblies, motivating the use of theoretical and simulation methods. This work advances a computational framework for conducting simulations of LLPS with residue-level detail, and allows for the determination of phase diagrams and coexistence densities of proteins in the two phases. The model includes a short-range contact potential as well as a simplified treatment of electrostatic energy. Interaction parameters are optimized against experimentally determined radius of gyration data for multiple unfolded or intrinsically disordered proteins (IDPs). These models are applied to two systems which undergo LLPS: the low complexity domain of the RNA-binding protein FUS and the DEAD-box helicase protein LAF-1. We develop a novel simulation method to determine thermodynamic phase diagrams as a function of the total protein concentration and temperature. We show that the model is capable of capturing qualitative changes in the phase diagram due to phosphomimetic mutations of FUS and to the presence or absence of the large folded domain in LAF-1. We also explore the effects of chain-length, or multivalency, on the phase diagram, and obtain results consistent with Flory-Huggins theory for polymers. Most importantly, the methodology presented here is flexible so that it can be easily extended to other pair potentials, be used with other enhanced sampling methods, and may incorporate additional features for biological systems of interest.

**Author summary:** Liquid liquid phase separation (LLPS) of low-complexity protein sequences has emerged as an important research topic due to its relevance to membraneless organelles and intracellular compartmentalization. However a molecular level understanding of LLPS cannot be easily obtained by experimental methods due to difficulty of determining structural properties of phase separated protein assemblies, and of choosing appropriate mutations. Here we advance a coarse-grained computational framework for accessing the long time scale phase separation process and for obtaining molecular details of LLPS, in conjunction with state of the art enhanced sampling methods. We are able to capture qualitatively the changes of phase diagram due to specific mutations, inclusion of a folded domain, and to variation of chain length. The model is flexible and can be used with different knowledge-based potential energy functions, as we demonstrate. We expect a wide application of the presented framework for advancing our understanding of the formation of liquid-like protein assemblies.

## Introduction

Intracellular compartmentalization is essential for normal physiological activity. This is commonly accomplished through isolation by lipid membranes or vesicles, but can also be achieved without the use of a membrane via membraneless organelles [1–3]. These organelles include processing bodies [4], stress granules [3, 5–7] and germ granules [8, 9] in the cytoplasm, and nucleoli [10] and nuclear speckles [11] in the nucleus. It has been recently established that many of these membraneless organelles can be described as phase separated liquid-like droplets [8, 12]. The process of liquid-liquid phase separation (LLPS) allows these organelles to spontaneously coalesce and disperse, and is important for many biological functions, such as response to heat shock and other forms of stress [6, 13, 14], DNA repair [15, 16], regulation of gene expression [17, 18], cellular signaling [3, 19], and many other functions requiring spatial organization and biochemical regulation [10, 20–22]. LLPS has also been implicated as a precursor to the formation of hydrogels [23] and fibrillar aggregates [7, 15], suggesting possible relevance to the pathogenesis of many diseases including Amyotrophic Lateral Sclerosis (ALS) and Frontotemporal Dementia (FTD) [15, 24].

Experimental studies have characterized different properties of biological LLPS, and have shown that many of these assemblies share several common characteristics. First, the formation and dissolution processes can be tuned by the cellular environment such as changes in temperature, pH and salt concentration [25], by post-translational modification such as phosphorylation [19, 26], and by mixing with other biomolecules such as proteins [27], RNA [28–30], and ATP [2, 30]. Second, the concentrated phase has liquid-like properties, including fusion, dripping, wetting [25] and ostwald ripening [28]. Its viscosity is typically several orders of magnitude higher than that of water [2, 8, 25]. Third, the LLPS is commonly driven or modulated by low complexity (LC) intrinsically disordered regions (IDRs) of the protein sequence [6, 25, 31], suggesting similarities to the well-characterized LLPS of polymer mixtures [32]. It should be noted that a disordered domain is not necessary for LLPS to occur [14], and indeed LLPS is known to occur for folded proteins during crystallization or purification [33]. Folded domains along with IDRs have also been shown to modulate LLPS properties [34]. Lastly, some proteins involved in LLPS process are also able to form fibril structure [7, 15], suggesting a possible connection between the liquid-like droplet and solid-like fibril states. However, the molecular level understanding of LLPS cannot be easily obtained by experimental methods due to difficulty of obtaining structural properties even in the concentrated phase [6], and the cumbersome process of screening mutations [35].

A number of recent theoretical and simulation studies have addressed protein phase separation. Jacobs and Frenkel used Monte Carlo simulations to study multiple-component phase separation and found that the phase boundary is very sensitive to intermolecular interactions, but less dependent on the number of components in the system [36]. Lin and Chan applied the random phase approximation to treat electrostatic interactions [37] and Flory Huggins theory for mixing entropy and other interactions. They were able to capture the sequence specificity of charged amino acids and found that the dependence of the phase boundary of the IDP Ddx4 on salt concentration can be explained by considering only electrostatic screening in their model [38]. It was also found that the monomer radius of gyration (*R_g_*) is correlated with the corresponding critical temperature in both theoretical work [39] and by experiment [14]. This supports the hypothesis that fundamental polymer physics principles can be used to understand LLPS [40]. However, a computational framework capable of capturing the general sequence specificity including both hydrophobic and electrostatic interactions [35] and molecular details on both intra- and inter-molecular interactions is still missing. All-atom simulation has the potential of fulfilling both tasks [41, 42] with the use of force fields suitable for intrinsically disordered proteins (IDPs) [43, 44]. Such a force field has been recently applied to study the monomer properties of TDP-43 which is known to undergo LLPS [45]. However, computational efficiency imposes limits on the use of all-atom representation for simulating LLPS directly. Even the use of coarse-grained simulations requires well-designed sampling methods to overcome the enthalpy gap of the first order phase transition [46, 47].

In this work, we introduce a general computational framework for studying LLPS, combining a residue based potential capable of capturing the sequence specific interactions and the slab simulation method capable of achieving convergence for phase transition properties including critical temperature, and the protein concentrations in dilute and concentrated phases. To demonstrate the capabilities of the model, we have selected two model proteins: the LC domain of RNA-binding protein Fused in Sarcoma (FUS), and DEAD-box helicase protein LAF-1, both of which are able to phase separate in vitro and in vivo [6, 15, 25]. Mutations of FUS have been shown to be highly relevant to the pathogenesis of ALS [48, 49] and display the ability to alter the kinetics of both droplet formation and aggregation into fibrils [15]. In addition, both the full length and disordered domain of LAF-1 have been shown to phase separate in vitro [25], allowing us to explore the impact of a large, rigid, domain on the LLPS behavior, and how this framework performs for such systems.

The manuscript is organized as follows. First, we introduce our computational framework including the coarse-grained potential, the sampling method and the approach of considering folded proteins in the simulation. We then present the application of the method to two imporant systems. We first show the comparison of phase diagrams for wild-type (WT) FUS and a mutant, qualitatively consistent with recent experimental measurements. Secondly we demonstrate how inclusion of the folded domain alters the LAF-1 phase diagram. In both FUS and LAF-1, we show the flexibility of the framework by providing the results for two different coarse-grained potentials. Lastly, we investigate the phase diagram dependence on chain length, closely connect to the “multivalency” effect often discussed in connection with LLPS.

## Methods

### Coarse-grained Model Development

All-atom simulations are unable to reach the time scales needed to study phase separation with current state-of-the-art computational hardware resources and sampling methods. We therefore introduce a coarse-grained representation of the protein, in which each residue is represented as a single particle (Fig. 1a). The model takes into account the side-chain charges and chemical properties of the 20 different types of amino acids, listed in Table S1, thus making it sequence specific. The potential energy function contains bonded, electrostatic, and short-range pairwise interaction terms. Bonded interactions are modelled by a harmonic potential with a spring constant of 10 kJ/Å^2^ and a bond length of 3.8 A. Electrostatic interactions are modeled using a Coulombic term with Debye-Hückel [50] electrostatic screening to account for salt concentration, having the functional form:

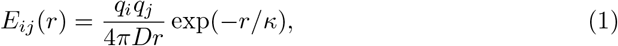

in which *κ* is the Debye screening length and *D* = 80, the dielectric constant of the solvent medium (water). For all the simulations for which phase diagrams are generated, a Debye screening length of 1 nm, corresponding to an ionic strength of approximately 100 mM, is used. For the IDP simulations to obtain *R_g_*, the ionic strength matching the experimental condition (Table S2) is used. The short-range pairwise potential accounts for both protein-protein and protein-solvent interactions. Here we have introduced two different models: the first is based on the amino acid hydrophobicity [51] and uses functional form introduced by Ashbaugh and Hatch [52]; the second is based on the Miyazawa-Jerningan potential [53] with the parameterized functional form taken from Kim and Hummer [54].

**Fig 1.**
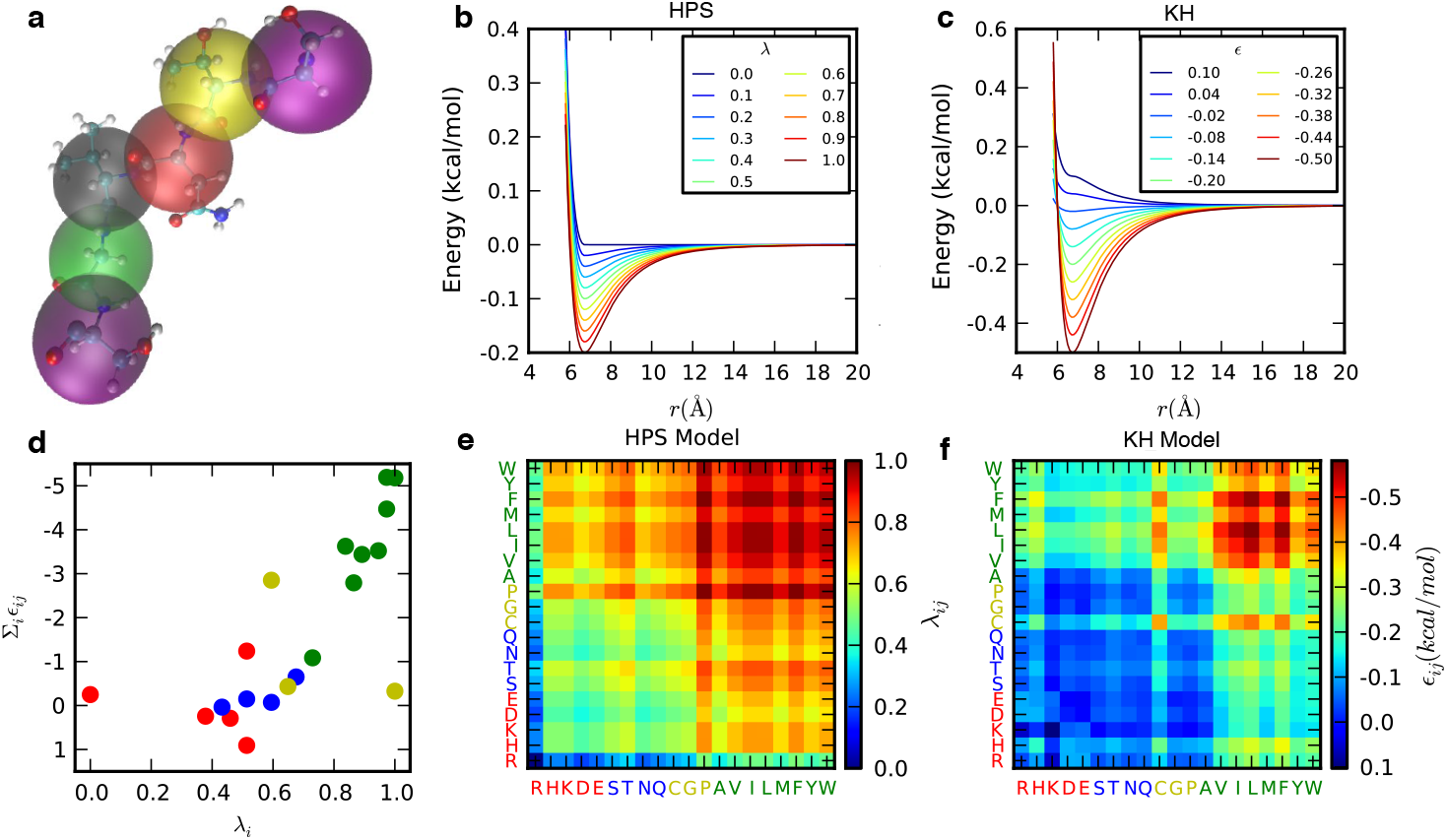
Schematic of the two knowledge-based potentials used for short-range pairwise interactions. a) Each amino acid is treated as a single particle. b, c) Potential energy functional form for HPS and KH models at different interaction strengths, shown with a constant σ value of 6 A. d) Correlation between the amino acid interaction strength (Σ_*i*_*ϵ_ij_*) in KH model and hydrophobicity (*λ_i_*) in HPS model, colored by the side-chain properties of amino acids (i.e., red for charged, blue for polar, green for hydrophobic and yellow for other amino acids). e, f) The pairwise interaction parameters used in HPS and KH model shown in color maps with blue being most repulsive interactions and red being most attractive. Amino acids are colored and clustered as described in d).

#### Hydrophobicity scale (HPS) model

The first model uses a hydrophobicity scale from the literature [51] to describe the effective interactions between amino acids. For use in the coarse-grained model, the atomic scale is first summed up to obtain a residue scale and is then scaled to the range from 0 to 1. The hydrophobicity values λ used for the 20 amino acids can be found in Table S1. The arithmetic average is set as the combination rule for both the pair interactions *λ* between two amino acids and the size σ of the amino acids (i.e., hydrophobicity scale *λ_ij_* = (*λ_i_* + *λ_j_*)/2 and amino acid size *σ_ij_* = (*σ_i_* + *σ_j_*)/2). The combined pairwise interaction strengths for each amino acid pair are shown in Fig. 1e. The Ashbaugh-Hatch functional form [52] which has previously been applied to the study of disordered proteins [55], allows the attractiveness of the interactions to be scaled by *λ* (Fig. 1b), and is described by,

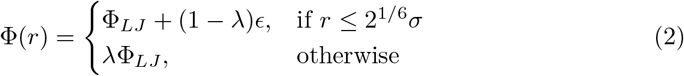

in which Φ_*LJ*_ is the standard Lennard-Jones potential

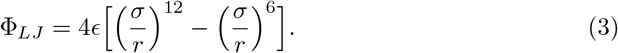

The pair potential for the least hydrophobic amino acid at a *λ* value of 0 consists of only the repulsive term, making it equivalent to the Weeks-Chandler-Andersen functional form [56]. The model contains one free parameter *ϵ*, which determines the absolute energy scale of the short-ranged interactions and is set to be constant across all pairs. To determine the optimal *ϵ*, *R_g_* was calculated for a set of IDPs (Table S2) using our model, and compared with available experimental *R_g_* data. Obtaining accurate estimates of R*g* from FRET and SAXS experimental data requires some care, as has recently been noted [57–60]. Since FRET probes an intramolecular pair distance, inferring *R_g_* requires the assumption of an underlying polymer model with known pair distance distribution and related *R_g_*. It has been shown that the commonly used Gaussian chain model works reasonably well for IDPs in the absence of chemical denaturants, but it breaks down when such denaturants are added [57, 61]. This is because the polymer scaling exponent *ν* ≈ 1/2 for IDPs without denaturants present, so that the Gaussian chain is a reasonable approximation for the denaturant-free conditions we are concerned with. We obtain the *R_g_* using a Gaussian chain model with a dye correction of 9 residues, as previously described [58, 62]. For SAXS, Guinier analysis is challenging because the approximation is only valid for a small range of q where the data tends to be noisy; when fitting a larger range of scatter angles, it tends to underestimate the *R_g_* [57]. A proper treatment of SAXS data requires a model that can also fit data at wider angles [57–59, 63]. Despite the limitations of the presently used data set, we expect that the systematic errors introduced by data analysis methods are still substantially smaller than the the deviation of the fit from experiment. However, a finer optimization of the model may require both the FRET and SAXS experimental data to be more accurately analyzed.

Fig. 2 and Fig. S1 show that an *ϵ* of 0.2 gives the greatest similarity to the experimental size of these unfolded proteins. In order to test if the model can capture the degree of collapse for folded and disordered sequences, we generated 131 random sequences with properties covering a wide range of net charge and hydrophobicity scale and determined their *R_g_* from simulation. In Fig. 3, we show that the *R_g_* of these sequences in a Uversky type plot [64] (i.e., as a function of the mean net charge 〈*q*〉 and the mean hydrophobicity 〈*h*〉 of the sequence) and in a Pappu type plot [65] (i.e., as a function of the fractions of the positive and negative charges in the sequence), both of which have been widely used to characterize sequence properties of proteins. *R_g_* values are observed ranging from 1.5 to 6.0 nm, and the predictions are good for naturally occurring test sequences. The larger *R_g_* values obtained for some of the synthetic sequences are outside the range observed for natural sequences in Fig. 2, however this is because the extreme synthetic sequences are essentially polyelectrolytes which are rare in nature. Although we do not have experimental data for such sequences, we note that the model still makes accurate predictions for the most charged protein in our data set, Prothymosin *α*-N (Table S2c), which has a net charge of −43 (−0.384 per residue), mean hydrophobicity of 0.555 and *R_g_* of 2.87 nm.

**Fig 2.**
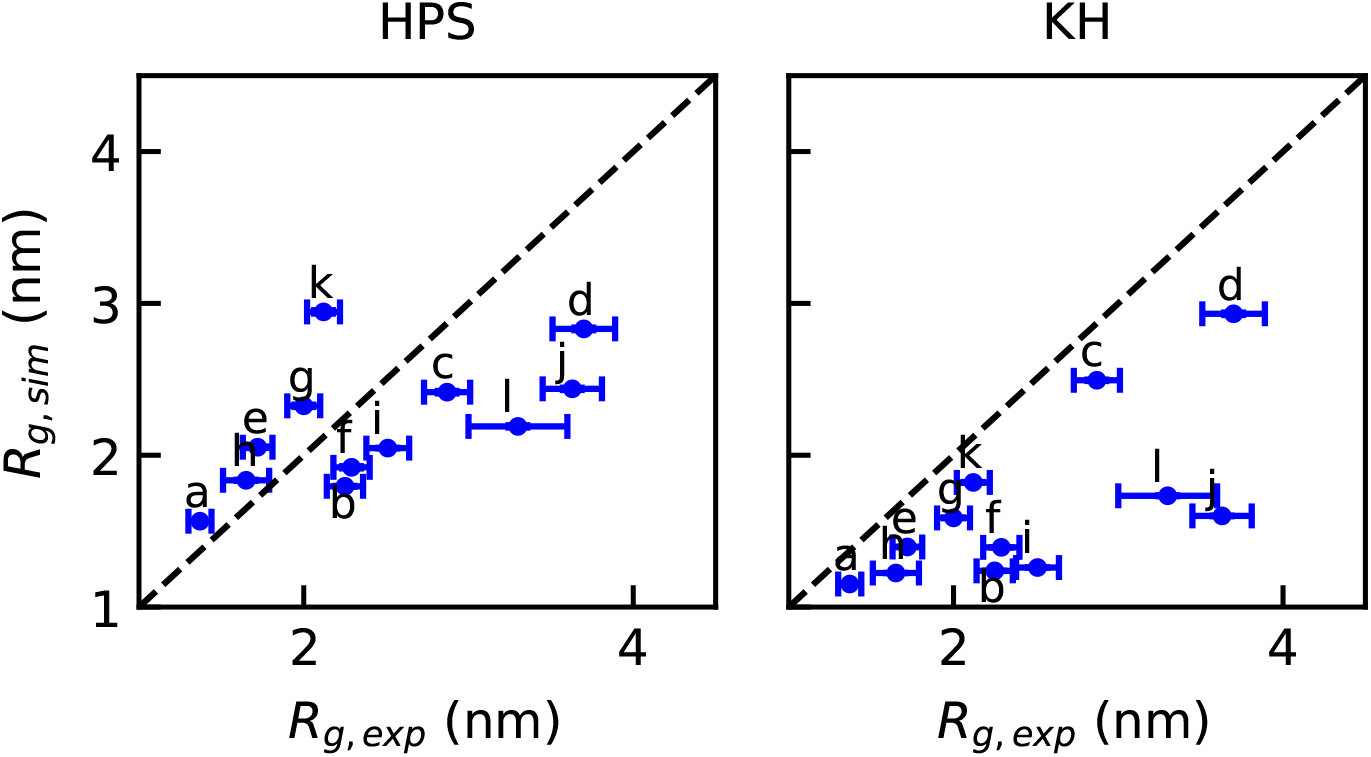
Parameterization of coarse-grained models: Comparison between radius of gyration of various intrinsically disordered proteins from experiment, and from simulation with the optimal parameters.

It is clear that the HPS model describes the known sequence-specific features of the disordered proteins, that is, a small mean hydrophobicity scale and a large mean net charge. The Uversky plot in Fig. 3 shows a correlation of *R_g_* with both hydrophobicity and mean charge per residue as seen in experiment [62]. It does appear that the correlation is stronger with net charge, while both factors were correlated with scaling exponents in earlier work [62]. This is partly because our sampled sequences span a larger range of charge, and also because charge and hydrophobicity are correlated in naturally occurring sequences, making it harder to separate their respective contributions. Even so, however, the correlation with charge does appear to be better also in experiment [62].

**Fig 3.**
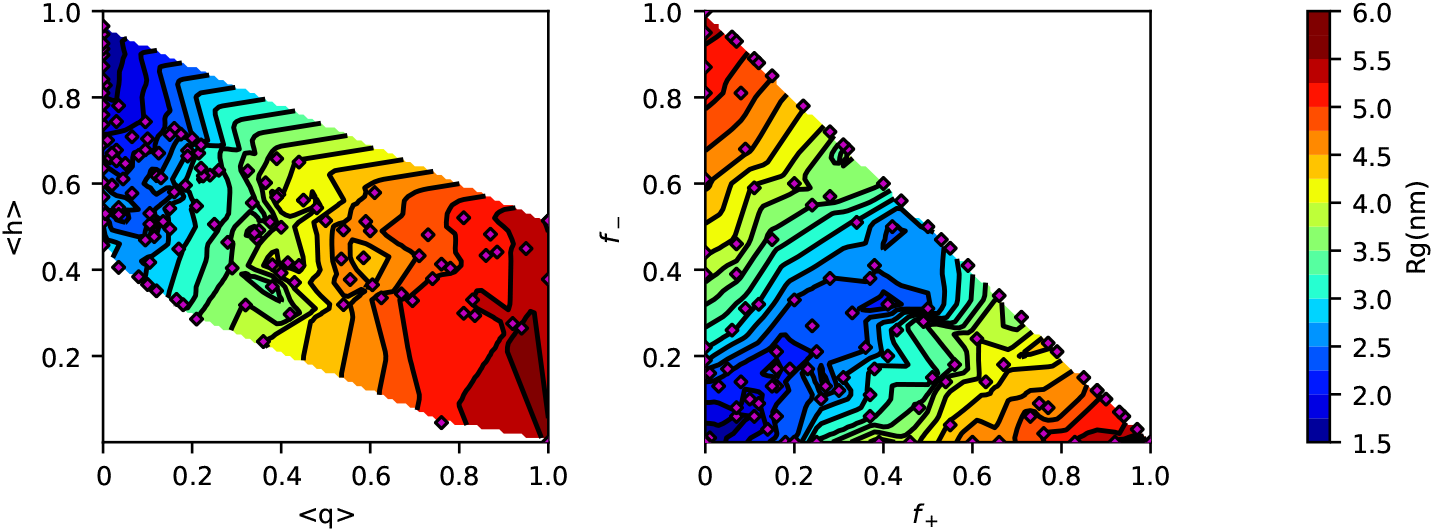
Randomly generated 100 amino acid sequences follow the general trends expected from an Uversky type plot (left) and Pappu type plot (right). Axes are: mean hydrophobicity per residue 〈*h*〉, mean net charge per residue, 〈*q*〉 and fractions of positively *f*_+_ and negatively *f*_−_ charged residues. For both plots, the color represents average *R_g_*, and contour lines are spaced every 0.25 nm. The location of each tested sequence is represented by a purple diamond.

#### Kim-Hummer (KH) model

A different model for short-range interactions has been previously developed and parameterized by Kim and Hummer to describe protein-protein interactions, using a variety of experimental data including the osmotic second virial coefficient of lysozyme and the binding affinity of the ubiquitin–CUE complex [54]. The potential function they used can be expressed in terms of Ashbaugh-Hatch potential function (Eq. 2 and 3), where

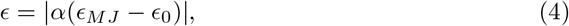

and

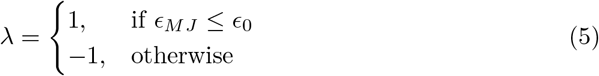

*ϵ_MJ_* is from the Miyazawa-Jerningan statistical contact potential [53]. Regarding the choice of *α* and *ϵ*_0_, the original literature identifies six sets of parameters, differing in the treatment of interactions involving buried residues. Here we employ parameter set *D* (*α* = 0.228 and *ϵ*_0_ = −1.00 kcal/mol, Table S4) for IDR, which generates a reasonable estimate of *R_g_* for a list of IDPs (Fig. 2), and parameter set A (*α* = 0.159 and *ϵ*_0_ = −1.36 kcal/mol, Table S5) for the helicase domain, which was parameterized for interactions between folded proteins [54], The correlation between the parameters of the HPS and KH models for IDR is shown in Fig. 1d. We repeat the analysis previously done with the HPS model on the same set of 100mers (Fig. S2) to provide additional insight into how the two models compare with regard to relative interaction strength of hydrophobic and electrostatic interactions. Both hydrophobic attraction and repulsion are stronger in the KH model than in HPS, and in this model there is a stronger dependence of the radius of gyration on hydrophobicity for sequences with low charge.

### Simulation framework

#### Slab method

In order to determine the phase diagram of the disordered proteins, we utilize a method [46, 66], in which the high-density (concentrated) phase, with surfaces normal to *z*, is simulated in equilibrium with the low-density (dilute) phase as shown in figure 4c and in supplementary movie S1. This allows the determination of the equilibrium density (or concentration) of proteins in each phase, and consequently the critical temperature, as described in more detail below. The initial configuration for the slab is prepared using the NPT ensemble with 100 protein chains at low temperature and high pressure to collapse them into a concentrated phase. This initial equilibration is conducted for 100 ns with periodic boundary conditions using a constant temperature (150 K), which is maintained by a Langevin thermostat with a friction coefficient of 1 ps^−1^ and pressure of 1 bar with a Parrinello-Rahman barostat [67]. A time step of 10 fs is used for all the simulations. The box size is first scaled to about 15 nm (25 nm for full length LAF-1) for both *x* and *y* axes and then equilibrated along the *z*-axis using anisotropic pressure coupling. Depending on the protein of interest and the pairwise potential functions, the length of the *z*-axis can vary. The *x*- and *y*- dimensions were set to 15 nm which is sufficient to prevent to most of the chains (> 99% estimated by a random-coil model for a 170-residue chain) from interacting with its periodic image. Then the *z*-dimension of the box was extended to 280 nm (~ 20 times larger than the initial z-dimension box size). Simulations are then run at multiple temperatures for ~ 5 *μ*s using constant temperature and volume with a Langevin thermostat. The temperature is gradually increased from 150 K to the targeted temperature over the first 100 ns. The next 1 *μ*s simulation is discarded as equilibration, and the remainder (at least 4 μs) is used for further analysis. The LAMMPS [68] and HOOMD-Blue v2.1.5 [69] software packages are used for molecular dynamics simulations in order to benefit from both CPU and GPU resources.

**Fig 4.**
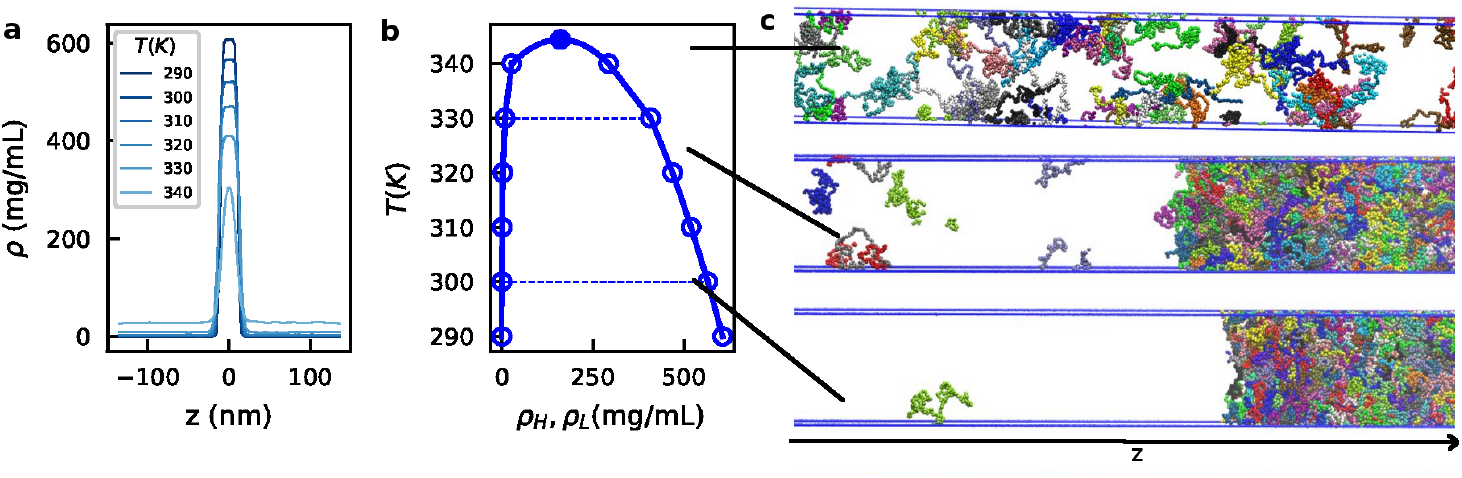
Slab sampling method and typical configurations in different regions of the phase diagram for FUS WT. a) Unsymmetrized slab density profile of FUS WT at different temperatures. The raw density plot is highly symmetrical, suggesting good convergence of simulations especially in converging the bulk density. b) Phase diagram of FUS WT obtained from the density profile. c) Typical configurations of FUS slab simulations at different temperatures.

We took several measures to verify that the initial configuration, system size and number of steps are sufficient to obtain well-converged thermodynamic properties of the system. First, we find that a simulation starting from a fully dispersed configuration, in which chains are put far from each other, but having the same periodic box geometry, will eventually coalesce to form a concentrated phase and generate a similar density profile (after 4*μ*s) to a simulation starting from a slab-like initial configuration (Fig. S3). Therefore a slab-like initial configuration reduces the length of the simulation required for convergence. Second, we do not see a quantitative difference of the results between the two halves of a 10*μ*s simulation (Fig. S4), suggesting 5 *μ*s is sufficient for convergence of the system. Third, we have also found that a system with 100 chains is sufficiently large to avoid finite-size effects, as the results are identical to those from a similar set of simulations containing 200 chains (Fig. S5).

#### Slab density profile

To determine the density profile along *z*, we first center the trajectory on the slab for each frame. The slab is defined as the cluster with the largest number of chains, clustering according to center-of-mass-distance between chain pairs. Chains with center-of-mass distances less than 5 nm are considered to be in the same cluster except for full length LAF-1, with which we use a cutoff of 7 nm due to its larger size. The density profile along *z* is then generated as shown in Fig. 4a and Fig. S6a. If phase separation occurs, we obtain the protein concentration of the dilute and concentrated phases (*ρ_L_* or *ρ_H_*) by using the average concentrations when |*z*| >50 nm or |*z*| <5 nm respectively. The protein concentration is shown in unit of mg/mL.

#### Phase diagram

The critical temperature *T_c_* can be obtained by fitting

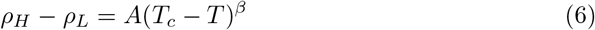

in which the critical exponent *β*=0.325 (universality class of 3D Ising model [70]) and A is a protein-specific fitting parameter. For fitting to this equation, a specific range of temperatures must be used. The minimum fitting temperature, *T*_1_, is chosen as the lowest temperature where *ρ_L_* is nonzero. The maximum fitting temperature *T*_2_ must be below the critical temperature as Eq. 6 can only describe the behavior below *T_c_* (Fig. S6c). To determine the optimal value for *T*_2_ we calculate the relative error of *T* when fitting *T* as a function of *ρ_H_* − *ρ_L_* using different test values of *T*_2_. This error will be large if *T*_2_ is greater than *T_c_* (Fig. S6d). We can then obtain a typical phase diagram as shown in Fig. 4b and Fig. S6b, in which the *ρ_L_* and *ρ_H_* when *T* < *T_c_* are determined from averaging different regions of the slab density profile as described above and *T_c_* and the corresponding *ρ_c_* are from fitting Eq. 6 (Fig. S6c). Fig. 4c shows visualizations of the different states of coexistence captured by these simulations. When the system is above *T_c_*, the slab evaporates to a supercritical protein solution. When the temperature is below *T_c_*, we see coexistence of two phases: one phase with free monomers and the other with many proteins in a condensed, liquid-like assembly. The number of free monomers decreases with decreasing temperatures to concentrations comparable with protein concentration in the dilute phase observed by experiment [6, 25].

#### Simulations with folded domain

Proteins which undergo LLPS usually contain multiple domains, including both folded and disordered domains [71]. Recently, Riback et al. found that poly(A)-binding protein Pab1 exhibits LLPS behavior in the absence of its disordered domain, but does not in the absence of the folded domains [14], contrary to the notion that intrinsic disorder is necessary for phase separation. Since both intrinsically disordered and folded domains can form favorable intermolecular interactions stabilizing the high density phase, it is only natural that they may both contribute to the LLPS behavior, and the contributions may be distributed differently from protein to protein. Therefore it would be beneficial to also introduce a framework capable of simulating LLPS containing rigid domains, using full-length LAF-1 which contains a folded domain and two disordered domains, as a test case.

The structure of the folded domain (helicase) of LAF-1 has not yet been solved, so we have used homology modelling and the Modeller v9.17 package [72] to embed the LAF-1 helicase sequence into its homologue with a solved crystal structure, VASA [73] (Fig. S7). Here we employ the KH model with parameter set A (*α* = 0.159 and *ϵ*_0_ = −1.36 kcal/mol) for all interactions involving the helicase domain, and parameter set D for interactions between disordered regions. The reason for this is that a 12-6 potential allows buried residues to make a significant contribution to binding energies of folded domains; this affects the affinity more than the specificity of the interactions. Model *A* was parameterized including such interactions for folded proteins, and is therefore appropriate for use in our model in describing interactions involving folded proteins. Model *D* was parameterized using a screening term to reduce the effect of buried residues, and is therefore appropriate for describing interactions between disordered regions where all residues are essentially fully exposed. A universal set of parameters would require a different functional form for the intermolecular interactions than the one we use here. When the structure of the folded domain is modelled, we treat the helicase as a rigid body (i.e., “fix rigid” command in LAMMPS or “md.constrain.rigid” command in HOOMD-Blue) in the simulation so that the structure of the folded domain is preserved. Interactions between residues within the same rigid body are neglected. The mass of the rigid body is scaled to be 0.5% of the original mass in order to accelerate rigid body dynamics. When calculating the density of the folded domain, the mass is scaled back to match the mass of the original folded domain with all residues. The folded domain can in principle also be simulated using harmonic restraints instead of rigid constraints, which would allow additional flexibility. However there is a clear advantage for using rigid body dynamics in terms of computational efficiency.

## Results

### Phase separation of FUS and its phosphomimetic mutants

As a first application of our model to LLPS, we use the prion-like LC domain of the protein FUS (FUS-LC) which is sufficient to induce LLPS in vitro in the absence of other biomolecules [15]. FUS-LC is an ideal system to test our model as it is fully disordered and displays very low secondary structure content [6]. The sequence is largely uncharged, with only 2 anionic aspartate residues within its 163 amino acid sequence. To test for sequence-specific effects, we conducted simulations for several different variants of the FUS-LC peptide, wild-type and four phosphomimetic mutants where a set of the 12 naturally phosphorylated threonine or serine residues are mutated to glutamate. [74] The first of these mutants is the 12E mutant, which contains all 12 glutamate substitutions, and does not undergo LLPS under similar conditions to FUS WT [75]. We additionally test the 6E mutant reported in the same work [75], and two designed variations of the 6E mutant, termed 6E’ and 6E* which maximize and minimize, respectively, the clustering of charged residues within the sequence under the constraints of preserving the amino acid composition of 6E, and only mutating naturally occurring phosphorylation sites. [76]

Utilizing the slab method, we have determined the range of temperatures at which the simulated FUS chains contain both phases, and have calculated the coexistence curve using both the HPS and KH models. The concentration of the dilute phase gives the predicted critical/saturation concentration of the protein, the concentration above which it will begin to form droplets in solution. The concentration of the dilute phase is on the order of 0.1-10 mg/mL over the tested temperature range, consistent with typical concentrations used to observe phase separation of FUS WT in vitro [6] (~1-5 mg/mL). We find that the critical temperature differs between the two models for FUS WT. However the coexistence curves and the phase diagrams are qualitatively similar(Fig. S9a), as are the intermolecular contact maps (Fig. S11).

To evaluate the impact of the phosphomimetic mutations, we determine the phase diagram for FUS WT, 6E, 6E’, 6E*, and 12E using the HPS model (Fig. 5). The 12E mutant phase separates at a much lower temperature, with the critical temperature smaller than even the lowest temperature at which we can observe coexistence between two phases for FUS WT (due to the prohibitively small concentration of the low-density phase). This is consistent with the experimental observation that FUS 12E is unable to phase separate in contrast to FUS WT at similar conditions [75]. The 6E mutants all lie between the two extreme cases, and have nearly identical phase diagrams as each other. While the difference of just 6 amino acids results in a greatly altered phase-separation ability, the rearrangement of these mutations does not seem to induce much change at all. However, these mutations were done under very strict constraints which do not allow for a significant change in the degree of charge clustering. We also calculate the interchain contacts, defined as when two amino acids of different chains are within 2^1/6^*σ_ij_* of each other. There are no specific contacts formed in either of the cases (Fig. S11), suggesting that LLPS of FUS WT is not driven by a specific region within the protein sequence. However, when comparing the different 6E mutants at the same temperature, the degree to which different regions of the peptide interact are greatly affected (Fig. S12). This shows that despite having almost identical phase coexistence, the interactions involved in phase separation can vary.

**Fig 5.**
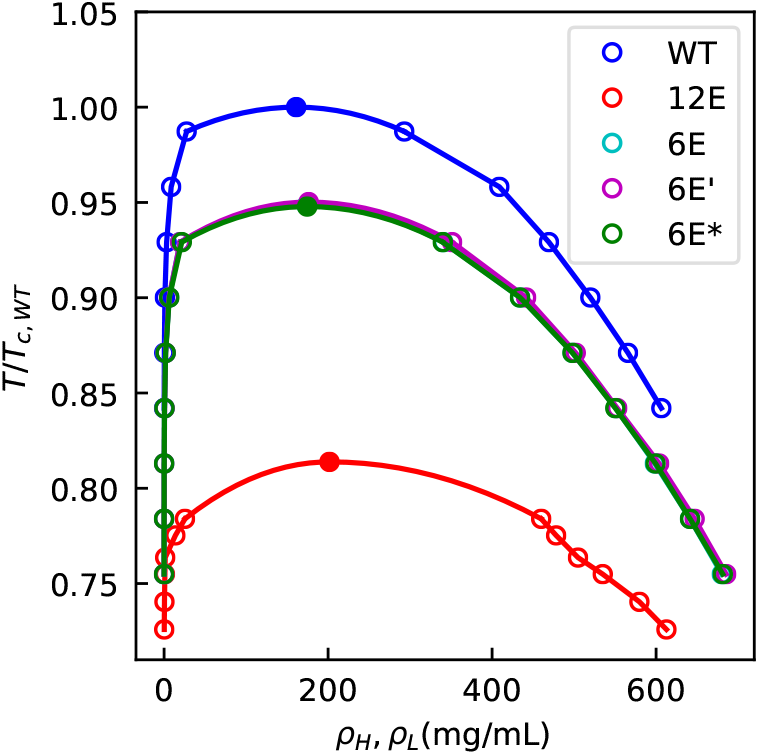
Phase diagram for FUS WT, 6E variants and 12E. Temperatures are scaled by the critical temperature of FUS WT.

To further check the liquid-like nature of the concentrated phase, we calculate the mean squared displacement (MSD) as a function of time for WT, 6E and 12E at 500mg/mL (Fig. S8, Supplementary Movie S2). For each, there is a linear region with non-zero slope suggesting that the concentrated phase is liquid-like, and not a solid aggregate. The diffusion coefficient from fitting the linear region is ~3×10^−6^ cm^2^/s, three orders of magnitude larger than measured in the experiment (4×10^−9^ cm^2^/s [6]) as can be expected from a coarse-grained simulation and the increased relaxation time used with Langevin dynamics. Finally, we check monomer radius of gyration in both the dilute and concentrated phase and find that chains in the concentrated phase are generally more extended than those in the dilute phase (Fig. S13).

### Phase separation of IDR and full length LAF-1

Next, we apply our model to DEAD-box helicase protein, LAF-1, which has been shown to phase separate as both its IDR and as full length, including a 437 residue folded domain, in vitro [25]. To test the effect of inclusion of folded domains, three variants of LAF-1 sequences have been simulated, including the N-terminal IDR of LAF-1, the helicase domain and its full length variant with both the IDR and folded domain as well as a short prion-like chain on the C-terminal. The IDR sequence contains a much higher fraction of charged amino acids (~26%) compared to FUS WT (~1%), and FUS 12E (~9%), and includes both attractive and repulsive electrostatic interactions. For IDR LAF-1, we simulated the phase diagram with both KH and HPS models. As was the case for FUS WT, the phase diagrams are qualitatively similar for the two models (Fig. S9b). For the helicase domain of LAF-1, the structure is predicted using homology modelling and kept rigid in simulations (details in the Methods section).

In Fig. 6 we compare the phase diagrams of the full-length and IDR regions of LAF-1. The phase diagram for the full length protein is shifted toward higher temperatures, and suggests a smaller saturation concentration as compared to the LAF-1 IDR alone at the same temperature. The results for the helicase domain alone also show clearly phase separation (Fig. S10). The experimental phase boundary in ~ 120 mM NaCl is ~ 0.05 mg/mL for full length LAF-1, but ~ 0.4 mg/mL for the isolated IDR [25]. Even though we cannot accurately estimate the low protein concentrations in the dilute phase so as to quantitatively compare with the experimental values, we do see an increase in the saturation concentration when adding the folded domain as has been seen by experiment. We note that the concentrations obtained from the high density phase are much higher than recently estimated by Wei et al. [77], however, they are quite comparable with those measured by Brady et al. for the similar DEAD-Box Helicase protein Ddx4 [78].

**Fig 6.**
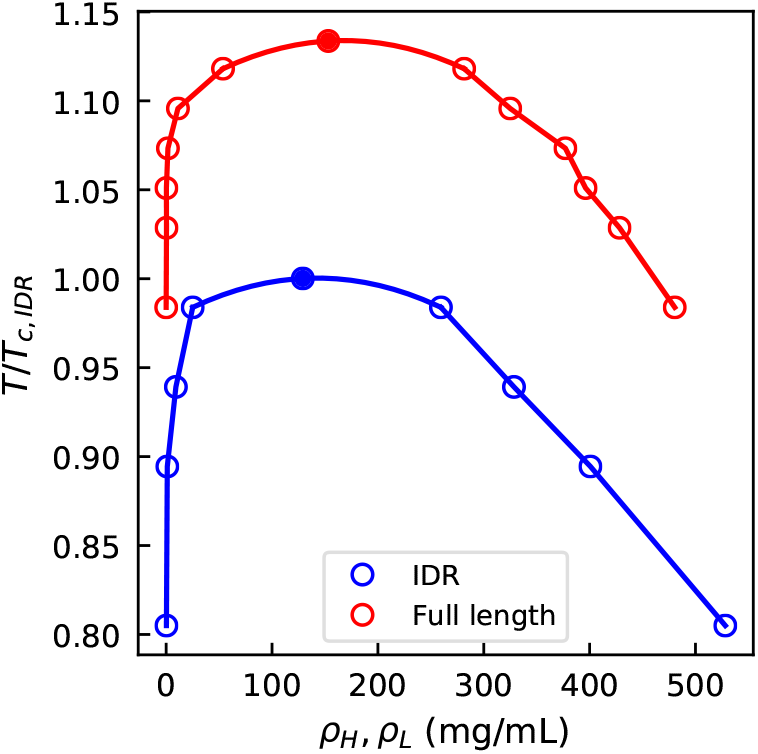
Phase diagram of IDR (blue) and full length (red) LAF-1. Temperatures are scaled by the critical temperature of IDR LAF-1. The corresponding slab density profiles are provided in Fig. S14.

The reason for the change of critical temperature upon inclusion of the folded domain is likely two-fold. First, the folded domain contains more hydrophobic residues with an average hydrophobicity (Table S1) of 0.664 (0.579 for the surface residues) in contrast to 0.520 for LAF-1 IDR, therefore strengthening the intermolecular attraction. In addition, providing more interaction sites per chain generally favors a higher critical temperature, because more interactions can be formed with a smaller loss of entropy, an effect usually referred to as multivalency [35]. The impact of multivalency on the critical temperature will be investigated explicitly in the next section.

In the concentrated phase, we also investigate the intermolecular contacts in Fig. 7. Unlike the case of FUS, there are regions along the sequence where there is a relatively high propensity to form contacts, (residue 21 to 28, RYVPPHLR) and (residue 13 to 18, NAALNR). These regions are present in both the IDR with the KH and HPS model (Fig. 7a and b) and in the full length protein (Fig. 7c). The central region of these two segments is composed of uncharged amino acids, suggesting the importance of hydrophobic patches in the sequence even with a large fraction of charged residues. As is shown in both 1D and 2D contact maps (Fig. 7a, c and d), the pattern of contacts and the number of contacts of the IDR look similar in the IDR and full length LAF-1 simulations. This observation also applies when comparing the contact maps between the helicase and full length LAF-1 simulations (Fig. S15). This suggests that the key residues contributing to the droplet formation are the same for the disordered peptide with and without the folded domain (Fig. 7d). Additionally, the disordered part of the protein (including both the N-terminal and C-terminal disordered regions) contributes more contacts than the folded domain in the simulation of full length LAF-1, consistent with the experimental observations that the disordered region of LAF-1 is the driving force for the LLPS [25]. The intramolecular contact map in the two phases (Fig. S16) supports the change of *R_g_* (Fig. S13) in that the peptide has fewer long range contacts in the concentrated phase than in the dilute phase.

**Fig 7.**
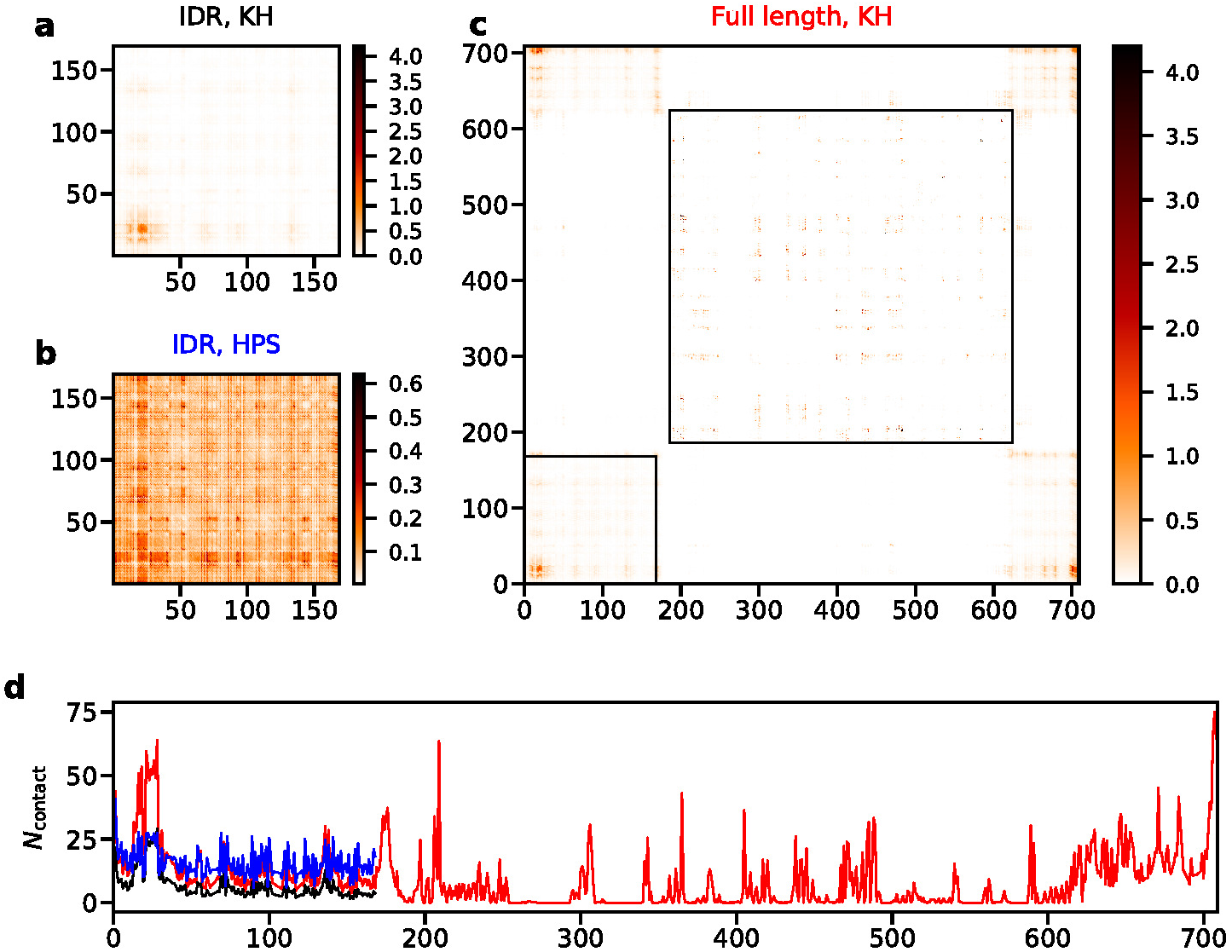
Number of intermolecular contacts per frame for LAF-1 with different models at 220K. a) Contact map of IDR LAF-1 with KH model. b) Contact map of IDR LAF-1 with HPS model. c) Contact map of full length LAF-1 with KH model. Black boxes illustrate the N-terminal IDR and the folded domain. d) Number of intermolecular contacts per residue per frame for IDR LAF-1 with KH model (black), IDR LAF-1 with HPS model (blue) and full length LAF-1 with KH model (red).

We additionally calculate the mean squared displacement (MSD) as a function of time for all the three variants of LAF-1 (i.e., IDR, helicase and full length) at concentrations predicted for the condensed phase at 210K (supplemental movies S4, S5 and S6), to see how the different regions affect the diffusion of the protein within the concentrated phase. There is a linear region with non-zero slope for all the variants (Fig. S8) suggesting liquid-like behavior. The IDR has a much larger diffusion coefficient than both the full length and the helicase domain of LAF-1 making it the most mobile of the three. This is likely due to its flexibility as well as the lower concentration. When comparing helicase with full length LAF-1, which are at similar concentrations, there is an order of magnitude difference in diffusion, further suggesting the importance of the flexible region of the sequence for maintaining the liquid-like behavior of proteins inside the droplet.

### Multivalency of IDRs

Multivalency has been shown to be important in driving LLPS in experiment [20, 35], in which proteins with more repeated units begin to form droplets at lower concentrations. Usually multivalency is used to describe a certain number of specific interaction sites per molecule. For polymers, there is inherently a large number of possible interactions between molecules, so for well-mixed sequences specific residue-residue interactions are less likely to play a role in assembly. Nonetheless, increasing the chain length will (for a given sequence composition) increase the number of available interaction sites per chain, and hence multivalency.

In order to investigate the mechanism of such behavior, we use a model system where we take the first 40 residues from the N-teriminus of FUS and make several repeated units in the form of [FUS40]_*n*_, in which n=1, 2, 3, 4 and 5. We then conduct multiple slab simulations for these repeated units of FUS fragment (see detailed system size in Table S3). In Fig. 8a we show the phase diagrams of [FUS40]_*n*_. It is clear that with increasing chain length, the phase boundary shifts to a higher temperature and a smaller concentration.

**Fig 8.**
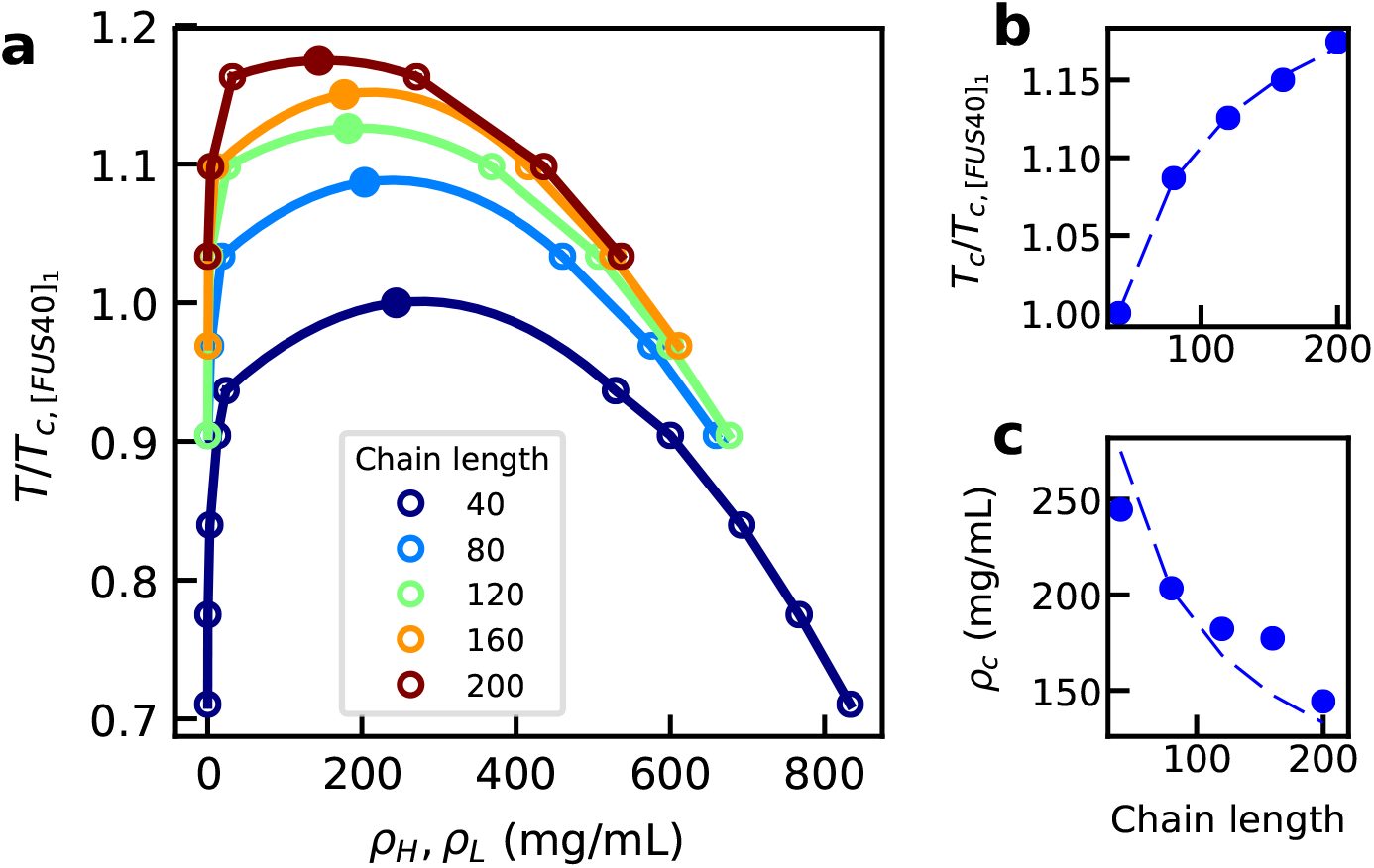
Phase separation of truncated FUS fragments of different lengths. a) Phase diagram for each peptide. b) The low critical temperature c) The critical concentration. Temperatures are scaled by the critical temperature of [FUS40]_1_. The corresponding slab density profiles are provided in Fig. S17.

To understand the mechanism of such dependence, we apply Flory-Huggins theory [79, 80], which has previously been used to understand IDP phase separation [9, 37, 38], to fit the phase transition properties obtained by molecular dynamics simulations when varying the chain length *N*. If we assume that each solvent molecule occupies one lattice position, then the the critical temperature 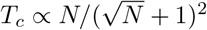 and the critical concentration 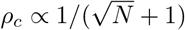. We can fit our simulated *T_c_* and *ρ_c_* as a function of the chain length with these approximating equations (assuming prefactor as the fitting parameter), as shown in Fig. 8b and c. These results suggest that the phase diagram dependence on the chain length can be described by Flory-Huggins theory. The term that is sensitive to the change of the chain length in this theory is the mixing entropy per segment. When increasing the chain length, the mixing entropy per segment decreases, and therefore the critical temperature increases. It would then be easier to observe LLPS with a longer chain at the same temperature, in the sense that the dilute-phase concentration is smaller, consistent with experimental observations [20]. This factor needs to be considered carefully when making mutations to protein sequences with the aim of understanding the molecular origin of LLPS: in general, chain truncation or extension will disfavor or favor LLPS, respectively, regardless of the sequence-specific effects. Similarly, when cutting a larger protein into fragments in order to evaluate the contribution of each to driving LLPS, it is expected in general that longer fragments will be able to phase separate at a higher temperature.

## Conclusion

We have introduced a general framework for conducting molecular dynamics simulations of LLPS leading to protein assemblies constituting many membraneless organelles. The level of coarse-graining, amino-acid-resolution, gives access to length and time scales needed to observe this phenomenon, and to achieve convergence of thermodynamic observables (i.e., phase diagram, critical temperature and protein concentration in the dilute and concentrated phases) while still allowing observance of changes induced by mutations to the protein sequence. The force fields utilized in this work are based on previously determined, knowledge-based potentials, parameterized to accurately represent radius of gyration of disordered proteins, but the framework is also flexible to incorporate other residue-based pairwise interaction potentials. The two force fields generate similar intermolecular contact maps within the concentrated phase, suggesting that description of the weak nonspecific interactions in IDPs can be captured easily with different models as compared to the description of specific interactions involving binding between folded proteins.

We have tested the framework and the two force fields with two model systems, which undergo phase separation in vitro, which yield phase diagrams giving the critical temperature, and saturation concentration at the tested temperatures. Despite that simplicity of the currently used potentials, and the fact that they were exclusively optimized based on the properties of monomeric proteins, we demonstrate the ability to predict how various perturbations to the system can change the LLPS. In the case of FUS LC, the model is able to capture the experimentally observed variation of phase diagram when introducing mutations. In LAF-1, the model is able to capture the experimentally observed difference between the phase separation of full length and truncated disordered-only sequences.

We have also investigated an important feature of LLPS regarding the dependence of phase behavior on chain length, which is well established in polymer physics and was previously observed in experiment [20]. We show that there is an upward shift in the phase diagram (temperature-concentration) with increasing chain length, with the critical point moving toward lower concentration and a higher temperature. At a given temperature, the saturation concentration will be higher for shorter chain lengths. Both the critical temperature and concentration are in good agreement with Flory-Huggins theory and therefore suggest the behavior can be explained by relative loss of entropy. With this in mind, if the phase behavior of interesting proteins cannot be observed in vitro, making repeated units might be a convenient way to shift the phase diagram toward being observable under experimental conditions. One must also consider this effect when making changes to protein length, such as His tags, or cleavage of a certain section of residues, and how just the change in chain length may affect the phase behavior.

Additionally, we measure certain important properties of proteins within the concentrated phase for the two model systems such as intermolecular contact propensities, which are quite difficult to resolve experimentally. With FUS LC, the intermolecular contacts are evenly distributed throughout the length of the peptide, suggesting that non-specific hydrophobic interactions are largely responsible for driving the phase-separation. For LAF-1, we observe excessive intermolecular contacts within a specific region (residue 21-29), largely composed of hydrophobic side-chains, suggesting that even though LAF-1 is 26% charged in sequence, hydrophobic interactions are still an important driving force for LLPS. The identification of this region with enhanced contact propensity shows that the model could serve as an initial screening tool for future mutagenesis studies.

There are some features that cannot be captured in the presented model, but can be added in the future work. First the absolute temperature of the simulation is not comparable to the experiment. The phase behavior at the lower critical solution temperature, which is observed in some disordered peptides experimentally [31], cannot be captured, either. Both require the addition of a temperature dependent solvation energy term into the framework, and more experimental *R_g_* data (or other relevant data) on IDPs to optimize the free parameters in the interaction potential. Second, we have not fully tested the ionic strength dependence data even though the trend is captured in LAF-1 with the current model, due to the breakdown of Debye-Hückle electrostatic screening at high ionic strength. However, we do not see any ionic strength dependence for FUS LC, which is inconsistent with the experiment [6]. To capture salt dependence in proteins with negligible charged amino acid content, it may be necessary to include a description of “salting-out” effects, i.e., the change of solubility with salt concentration as captured by the Hofmeister series. In Fig. S18, we show that the literature-known amino acid specific salting-out coefficients [81–84] are strongly correlated with the hydrophobicity scale and therefore it may be possible to model the salting-out effect with an additional energy term using the same hydrophobicity scale. In the future, one would also like to introduce additional handles (such as a structure-based potential for intramolecular interactions) to allow for conformational changes within the folded parts of a chain. This will allow one to study LLPS of proteins with small populations of folded regions that are important for self-assembly, such as transient helical regions in the disordered domain of TDP-43 [45].

## Supporting information

**Fig. S1** Comparison of *R_g_* between simulations and experiments with different e parameters for HPS model.

**Fig. S2** Pappu and Uversky plots for set of 100mers with different properties using KH-D model.

**Fig. S3** LAF-1 simulation started from dispersed state at 210K density profiles at different times showing coalescence to dense phase.

**Fig. S4** Comparison of density profiles between first 5*μ*s and last 5*μ*s of slab simulations of FUS WT.

**Fig. S5** Comparison of FUS WT simulations with 100 (blue) and 200 (red) chains.

**Fig. S6** Illustrations of the methodology to determine the range of temperatures to fit Eq. 7 in the main text.

**Fig. S7** Homology modelling of helicase domain of LAF-1 using the structure of VASA.

**Fig. S8** Mean squared displacement (MSD) as a function of time for FUS variants at 260K and LAF-1 variants variants at 220K.

**Fig. S9** Comparison of the phase diagram generated with HPS (blue) and KH (red) model in FUS WT (left) and LAF-1 IDR (right).

**Fig. S10** Phase diagram of IDR (blue), helicase (cyan) and full length (red) LAF-1. Temperatures are scaled by the critical temperature of IDR LAF-1.

**Fig. S11** Inter- (upper) and intra-molecular (lower) contact maps for FUS WT at 260 K using HPS (left) and KH models (right).

**Fig. S12** Intermolecular contacts for FUS 6E’ divided by that of FUS 6E* showing how the overall number of contacts forming within the slab changes between the two sequences.

**Fig. S13** Radii of gyration of the disordered proteins inside (blue) and out of (red) the slab.

**Fig. S14** Slab density profiles of IDR (left), helicase (middle) or full length (right) LAF-1.

**Fig. S15** Number of intermolecular contacts per frame for different LAF-1 variants at 220K. a) Contact map of IDR LAF-1. b) Contact map of helicase LAF-1. c) Contact map of full length LAF-1. Black boxes illustrate the N-terminal IDR and the helicase domain. d) Number of intermolecular contacts per residue per frame for IDR LAF-1 (black), helicase LAF-1 (blue) and full length LAF-1 (red).

**Fig. S16**Number of intramolecular contacts for LAF1 IDR with KH model.

**Fig. S17**Slab density profiles of the repeated peptides of FUS fragment.

**Fig. S18**The correlation between salting-out constant and hydrophobicity scale.

**Table S1** The amino acid parameters used in the HPS model.

**Table S2** List of intrinsically disordered or unfolded proteins with experimentally determined *R_g_*.

**Table S3** Summary of slab simulations and critical temperatures obtained.

**Table S4** Interaction parameters (*ϵ_ij_*) used for KH-D model.

**Table S5** Interaction parameters (*ϵ_ij_*) used for KH-A model.

## Acknowledgements

J.M. is thankful to Professor Thanos Panagiotopoulos for hospitality during his sabbatical stay at Princeton University. We thank Mike Howard (Princeton University) for useful discussions on the slab method and HOOMD, Prof. Nick Fawzi (Brown University) for discussions on FUS, and Prof. Cliff Brangwynne (Princeton University) for discussions on LAF-1. This research is supported by US Department of Energy (DOE), Office of Science, Basic Energy Sciences (BES) under Award DE-SC0013979. This research used resources of the National Energy Research Scientific Computing Center, a DOE Office of Science User Facility supported under Contract No. DE-AC02-05CH11231. Use of the high-performance computing capabilities of the Extreme Science and Engineering Discovery Environment (XSEDE), which is supported by the National Science Foundation, project no. TG-MCB120014, is also gratefully acknowledged. R.B. and W.Z. were supported by the intramural research program of the National Institute of Diabetes and Digestive and Kidney Diseases of the National Institutes of Health. This work utilized the computational resources of the NIH HPC Biowulf cluster. (http://hpc.nih.gov)

## References

1. An S, Kumar R, Sheets ED, Benkovic SJ. Reversible compartmentalization of de novo purine biosynthetic complexes in living cells. Science. 2008;320(5872):103–106.

2. Brangwynne CP, Mitchison TJ, Hyman AA. Active liquid-like behavior of nucleoli determines their size and shape in Xenopus laevis oocytes. Proc Natl Acad Sci USA. 2011;108(11):4334–4339.

3. Wippich F, Bodenmiller B, Trajkovska MG, Wanka S, Aebersold R, Pelkmans L. Dual specificity kinase DYRK3 couples stress granule condensation/dissolution to mTORC1 signaling. Cell. 2013;152(4):791–805.

4. Fromm SA, Kamenz J, Nöldeke ER, Neu A, Zocher G, Sprangers R. In Vitro Reconstitution of a Cellular Phase-Transition Process that Involves the mRNA Decapping Machinery. Angew Chem Int Edit. 2014;53(28):7354–7359.

5. Kato M, Han TW, Xie S, Shi K, Du X, Wu LC, et al. Cell-free formation of RNA granules: low complexity sequence domains form dynamic fibers within hydrogels. Cell. 2012;149(4):753–767.

6. Burke KA, Janke AM, Rhine CL, Fawzi NL. Residue-by-residue view of in vitro FUS granules that bind the C-terminal domain of RNA polymerase II. Mol Cell. 2015;60(2):231–241.

7. Molliex A, Temirov J, Lee J, Coughlin M, Kanagaraj AP, Kim HJ, et al. Phase separation by low complexity domains promotes stress granule assembly and drives pathological fibrillization. Cell. 2015;163(1):123–133.

8. Brangwynne CP, Eckmann CR, Courson DS, Rybarska A, Hoege C, Gharakhani J, et al. Germline P granules are liquid droplets that localize by controlled dissolution/condensation. Science. 2009;324(5935):1729–1732.

9. Nott TJ, Petsalaki E, Farber P, Jervis D, Fussner E, Plochowietz A, et al. Phase transition of a disordered nuage protein generates environmentally responsive membraneless organelles. Mol Cell. 2015;57(5):936–947.

10. Feric M, Vaidya N, Harmon TS, Mitrea DM, Zhu L, Richardson TM, et al. Coexisting liquid phases underlie nucleolar subcompartments. Cell. 2016;165(7):1686–1697.

11. Marzahn MR, Marada S, Lee J, Nourse A, Kenrick S, Zhao H, et al. Higher-order oligomerization promotes localization of SPOP to liquid nuclear speckles. EMBO J. 2016; p. e201593169.

12. Uversky VN. Protein intrinsic disorder-based liquid–liquid phase transitions in biological systems: Complex coacervates and membrane-less organelles. Adv Colloid Interfac. 2017;239:97–114.

13. Biamonti G, Vourc’h C. Nuclear stress bodies. Cold Spring Harb Perspect Biol. 2010;2(6):a000695.

14. Riback JA, Katanski CD, Kear-Scott JL, Pilipenko EV, Rojek AE, Sosnick TR, et al. Stress-triggered phase separation is an adaptive, evolutionarily tuned response. Cell. 2017;168(6):1028–1040.

15. Patel A, Lee HO, Jawerth L, Maharana S, Jahnel M, Hein MY, et al. A liquid-to-solid phase transition of the ALS protein FUS accelerated by disease mutation. Cell. 2015;162(5):1066–1077.

16. Altmeyer M, Neelsen KJ, Teloni F, Pozdnyakova I, Pellegrino S, Grøfte M, et al. Liquid demixing of intrinsically disordered proteins is seeded by poly (ADP-ribose). Nat Commun. 2015;6:8088.

17. Morimoto M, Boerkoel CF. The role of nuclear bodies in gene expression and disease. Biology. 2013;2(3):976–1033.

18. Hnisz D, Shrinivas K, Young RA, Chakraborty AK, Sharp PA. A phase separation model for transcriptional control. Cell. 2017;169(1):13–23.

19. Su X, Ditlev JA, Hui E, Xing W, Banjade S, Okrut J, et al. Phase separation of signaling molecules promotes T cell receptor signal transduction. Science. 2016;352(6285):595–599.

20. Li P, Banjade S, Cheng HC, Kim S, Chen B, Guo L, et al. Phase transitions in the assembly of multi-valent signaling proteins. Nature. 2012;483(7389):336.

21. Jiang H, Wang S, Huang Y, He X, Cui H, Zhu X, et al. Phase transition of spindle-associated protein regulate spindle apparatus assembly. Cell. 2015;163(1):108–122.

22. Nott TJ, Craggs TD, Baldwin AJ. Membraneless organelles can melt nucleic acid duplexes and act as biomolecular filters. Nat Chem. 2016;8(6):569–575.

23. Xiang S, Kato M, Wu LC, Lin Y, Ding M, Zhang Y, et al. The LC domain of hnRNPA2 adopts similar conformations in hydrogel polymers, liquid-like droplets, and nuclei. Cell. 2015;163(4):829–839.

24. Mateju D, Franzmann TM, Patel A, Kopach A, Boczek EE, Maharana S, et al. An aberrant phase transition of stress granules triggered by misfolded protein and prevented by chaperone function. EMBO J. 2017; p. e201695957.

25. Elbaum-Garfinkle S, Kim Y, Szczepaniak K, Chen CCH, Eckmann CR, Myong S, et al. The disordered P granule protein LAF-1 drives phase separation into droplets with tunable viscosity and dynamics. Proc Natl Acad Sci USA. 2015;112(23):7189–7194.

26. Wang JT, Smith J, Chen BC, Schmidt H, Rasoloson D, Paix A, et al. Regulation of RNA granule dynamics by phosphorylation of serine-rich, intrinsically disordered proteins in C. elegans. Elife. 2014;3:e04591.

27. Lin Y, Protter DS, Rosen MK, Parker R. Formation and maturation of phase-separated liquid droplets by RNA-binding proteins. Mol Cell. 2015;60(2):208–219.

28. Berry J, Weber SC, Vaidya N, Haataja M, Brangwynne CP. RNA transcription modulates phase transition-driven nuclear body assembly. Proc Natl Acad Sci USA. 2015;112(38):E5237–E5245.

29. Zhang H, Elbaum-Garfinkle S, Langdon EM, Taylor N, Occhipinti P, Bridges AA, et al. RNA controls PolyQ protein phase transitions. Mol Cell. 2015;60(2):220–230.

30. Kim Y, Myong S. RNA Remodeling Activity of DEAD Box Proteins Tuned by Protein Concentration, RNA Length, and ATP. Mol Cell. 2016;63(5):865–876.

31. Quiroz FG, Chilkoti A. Sequence heuristics to encode phase behaviour in intrinsically disordered protein polymers. Nat Mater. 2015;14(11):1164.

32. Bates FS. Polymer-polymer phase behavior. Science. 1991;251(4996):898–905.

33. Asherie N. Protein crystallization and phase diagrams. Methods. 2004;34(3):266–272.

34. Shin Y, Berry J, Pannucci N, Haataja MP, Toettcher JE, Brangwynne CP. Spatiotemporal control of intracellular phase transitions using light-activated optoDroplets. Cell. 2017;168(1):159–171.

35. Pak CW, Kosno M, Holehouse AS, Padrick SB, Mittal A, Ali R, et al. Sequence determinants of intracellular phase separation by complex coacervation of a disordered protein. Mol Cell. 2016;63(1):72–85.

36. Jacobs WM, Frenkel D. Phase transitions in biological systems with many components. Biophys J. 2017;112(4):683–691.

37. Lin YH, Song J, Forman-Kay JD, Chan HS. Random-phase-approximation theory for sequence-dependent, biologically functional liquid-liquid phase separation of intrinsically disordered proteins. J Mol Liq. 2017;228:176–193.

38. Lin YH, Forman-Kay JD, Chan HS. Sequence-specific polyampholyte phase separation in membraneless organelles. Phys Rev Lett. 2016;117(17):178101.

39. Lin YH, Chan HS. Phase Separation and Single-Chain Compactness of Charged Disordered Proteins Are Strongly Correlated. Biophys J. 2017;.

40. Brangwynne CP, Tompa P, Pappu RV. Polymer physics of intracellular phase transitions. Nat Phys. 2015;11(11):899.

41. Shaw DE, Maragakis P, Lindorff-Larsen K, Piana S, Dror RO, Eastwood MP, et al. Atomic-level characterization of the structural dynamics of proteins. Science. 2010;330:341–346.

42. Lindorff-Larsen K, Piana S, Dror RO, Shaw DE. How fast-folding proteins fold. Science. 2011;334:517–520.

43. Best RB, Zheng W, Mittal J. Balanced protein-water interactions improve properties of disordered proteins and non-specific protein association. J Chem Theor Comput. 2014;10:5113–5124.

44. Piana S, Donchev AG, Robustelli P, Shaw DE. Water dispersion interactions strongly influence simulated structural properties of disordered protein states. J Phys Chem B. 2015;119:5113–5123.

45. Conicella AE, Zerze GH, Mittal J, Fawzi NL. ALS mutations disrupt phase separation mediated by *α*-helical structure in the TDP-43 low-complexity C-terminal domain. Structure. 2016;24(9):1537–1549.

46. Blas FJ, MacDowell LG, de Miguel E, Jackson G. Vapor-liquid interfacial properties of fully flexible Lennard-Jones chains. J Chem Phys. 2008;129(14):144703.

47. Kim J, Keyes T, Straub JE. Generalized replica exchange method. J Chem Phys. 2010;132(22):224107.

48. Vance C, Rogelj B, Hortobágyi T, De Vos KJ, Nishimura AL, Sreedharan J, et al. Mutations in FUS, an RNA processing protein, cause familial amyotrophic lateral sclerosis type 6. Science. 2009;323(5918):1208–1211.

49. Kwiatkowski TJ, Bosco D, Leclerc A, Tamrazian E, Vanderburg C, Russ C, et al. Mutations in the FUS/TLS gene on chromosome 16 cause familial amyotrophic lateral sclerosis. Science. 2009;323(5918):1205–1208.

50. Debye P, Hückel E. De la theorie des electrolytes. I. abaissement du point de congelation et phenomenes associes. Physikalische Zeitschrift. 1923;24(9):185–206.

51. Kapcha LH, Rossky PJ. A simple atomic-level hydrophobicity scale reveals protein interfacial structure. J Mol Biol. 2014;426(2):484–498.

52. Ashbaugh HS, Hatch HW. Natively unfolded protein stability as a coil-to-globule transition in charge/hydropathy space. J Am Chem Soc. 2008;130(29):9536–9542.

53. Miyazawa S, Jernigan RL. Residue-residue potentials with a favourable contact pair term and an unfavourable high packing density term, for simulation and threading. J Mol Biol. 1996;256:623–644.

54. Kim YC, Hummer G. Coarse-grained models for simulation of multiprotein complexes: application to ubiquitin binding. J Mol Biol. 2008;375:1416–1433.

55. Miller CM, Kim YC, Mittal J. Protein composition determines the effect of crowding on the properties of disordered proteins. Biophys J. 2016;111(1):28–37.

56. Weeks JD, Chandler D, Andersen HC. Role of repulsive forces in determining the equilibrium structure of simple liquids. J Chem Phys. 1971;56:5237–5247.

57. Borgia A, Zheng W, Buholzer K, Borgia MB, Schuler A, Hofmann H, et al. Consistent view of polypeptide chain expansion in chemical denaturants from multiple experimental methods. J Am Chem Soc. 2016;138(36):11714–11726.

58. Zheng W, Borgia A, Buholzer K, Grishaev A, Schuler B, Best RB. Probing the action of chemical denaturant on an intrinsically disordered protein by simulation and experiment. J Am Chem Soc. 2016;138(36):11702–11713.

59. Fuertes G, Banterle N, Ruff KM, Chowdhury A, Mercadante D, Koehler C, et al. Decoupling of size and shape fluctuations in heteropolymeric sequences reconciles discrepancies in SAXS vs. FRET measurements. Proc Natl Acad Sci USA. 2017; p. 201704692.

60. Song J, Gomes GN, Shi T, Gradinaru CC, Chan HS. Conformational Heterogeneity and FRET Data Interpretation for Dimensions of Unfolded Proteins. Biophys J. 2017;113:1012–1024.

61. O’Brien EP, Morrison G, Brooks BR, Thirumalai D. How accurate are polymer models in the analysis of Förster resonance energy transfer experiments on proteins? J Chem Phys. 2012;130:124903.

62. Hofmann H, Soranno A, Borgia A, Gast K, Nettels D, Schuler B. Polymer scaling laws of unfolded and intrinsically disordered proteins quantified with single-molecule spectroscopy. Proc Natl Acad Sci USA. 2012;109:16155–16160.

63. Riback JA, Bowman MA, Zmyslowski AM, Knoverek CR, Jumper JM, Hinshaw JR, et al. Innovative scattering analysis shows that hydrophobic proteins are expanded in water. Science. 2017;358:238–241.

64. Uversky VN, Gillespie JR, Fink AL. Why are “natively unfolded” proteins unstructured under physiologic conditions? Proteins. 2000;41:415–427.

65. Mao AH, Crick SL, Vitalis A, Chicoine C, Pappu RV. Net charge per residue modulates conformational ensembles of intriniscally disordered proteins. Proc Natl Acad Sci USA. 2010;107:8183–8188.

66. Silmore KS, Howard MP, Panagiotopoulos AZ. Vapour–liquid phase equilibrium and surface tension of fully flexible Lennard–Jones chains. Mol Phys. 2017;115(3):320–327.

67. Parrinello M, Rahman A. Polymorphic transitions in single crystals: a new molecular dynamics method. J Appl Phys. 1981;52:7182–7190.

68. Plimpton S. Fast parallel algorithms for short-range molecular dynamics. J Comput Phys. 1995;117(1):1–19.

69. Anderson JA, Lorenz CD, Travesset A. General purpose molecular dynamics simulations fully implemented on graphics processing units. J Comput Phys. 2008;227(10):5342–5359.

70. Rowlinson JS, Widom B. Molecular theory of capillarity. Courier Corporation; 2013.

71. Uversky VN, Kuznetsova IM, Turoverov KK, Zaslavsky B. Intrinsically disordered proteins as crucial constituents of cellular aqueous two phase systems and coacervates. FEBS Lett. 2015;589(1):15–22.

72. Šali A, Blundell TL. Comparative protein modelling by satisfaction of spatial restraints. J Mol Biol. 1993;234(3):779–815.

73. Sengoku T, Nureki O, Nakamura A, Kobayashi S, Yokoyama S. Structural basis for RNA unwinding by the DEAD-box protein Drosophila Vasa. Cell. 2006;125(2):287–300.

74. Kwon I, Kato M, Xiang S, Wu L, Theodoropoulos P, Mirzaei H, et al. Phosphorylation-regulated binding of RNA polymerase II to fibrous polymers of low-complexity domains. Cell. 2013;155(5):1049–1060.

75. Monahan Z, Ryan VH, Janke AM, Burke KA, Zerze GH, O’Meally R, et al. Phosphorylation of FUS low-complexity domain disrupts phase separation, aggregation, and toxicity. EMBO J. 2017;doi:10.15252/embj.201696394.

76. Sawle L, Ghosh K. A theoretical method to compute sequence dependent configurational properties in charged polymers and proteins. The Journal of chemical physics. 2015;143(8):08B615_1.

77. Wei MT, Elbaum-Garfinkle S, Holehouse AS, Chen CCH, Feric M, Arnold CB, et al. Phase behaviour of disordered proteins underlying low density and high permeability of liquid organelles. Nature Chemistry. 2017;.

78. Brady JP, Farber PJ, Sekhar A, Lin YH, Huang R, Bah A, et al. Structural and hydrodynamic properties of an intrinsically disordered region of a germ cell-specific protein on phase separation. Proceedings of the National Academy of Sciences. 2017;114(39):E8194–E8203.

79. Flory PJ. Thermodynamics of high polymer solutions. J Chem Phys. 1942;10(1):51–61.

80. Huggins ML. Some Properties of Solutions of Long-chain Compounds. J Phys Chem. 1942;46(1):151–158.

81. Schrier EE, Schrier EB. The salting-out behavior of amides and its relation to the denaturation of proteins by salts. J Phys Chem. 1967;71(6):1851–1860.

82. Nandi PK, Robinson DR. Effects of salts on the free energy of the peptide group. J Am Chem Soc. 1972;94(4):1299–1308.

83. Nandi PK, Robinson DR. Effects of salts on the free energies of nonpolar groups in model peptides. J Am Chem Soc. 1972;94(4):1308–1315.

84. Baldwin RL. How Hofmeister ion interactions affect protein stability. Biophys J. 1996;71(4):2056–2063.

